# DepLink: an R Shiny app to systematically link genetic and pharmacologic dependencies of cancer

**DOI:** 10.1101/2022.09.26.509353

**Authors:** Tapsya Nayak, Li-Ju Wang, Michael Ning, Gabriela Rubannelsonkumar, Eric Jin, Siyuan Zheng, Peter J. Houghton, Yufei Huang, Yu-Chiao Chiu, Yidong Chen

**Author notes:** Correspondence to Yu-Chiao Chiu, Ph.D. and Yidong Chen, Ph.D. Tapsya Nayak and Li-Ju Wang contributed equally to this work.

## Abstract

Large-scale genetic and pharmacologic dependency maps are generated to reveal genetic vulnerabilities and drug sensitivities of cancer. However, user-friendly software is needed to systematically link such maps. Here we present DepLink, an R Shiny server to identify genetic and pharmacologic perturbations that induce similar effects on cell viability or molecular changes. DepLink integrates heterogeneous datasets of genome-wide CRISPR loss-of-function screens, high-throughput pharmacologic screens, and perturbation expression signatures. The datasets are systematically connected by four complementary modules tailored for different query scenarios. In summary, DepLink enables easy navigation, visualization, and linkage of rapidly evolving cancer dependency maps.

## Background

With revolutionizing techniques in high-throughput screening and genomic profiling, large-scale genetic and pharmacologic dependency maps have been generated by systematically perturbing genes or treating cancer cell lines with small compounds [1-4]. These rapidly growing cancer dependency maps, also known as DepMap, play a significant role in precision oncology by identifying and targeting the “Achilles heel” of cancer [5, 6]. However, user-friendly software is needed to systematically link and visualize such complex maps to investigate similar mechanisms behind perturbations triggered by genes or compounds. To address this issue, here we present the DepLink application that enables researchers to identify genetic (gene knockouts) and pharmacologic perturbations (drug treatments) that induce similar effects. DepLink was designed to answer two important questions in cancer biology using large-scale data resources: i) which drugs can be potential surrogates to knockout of a gene in cancer cells, and ii) which genes are potential targets or mechanisms of action of a drug.

## Results and discussion

As shown in Fig. 1A, DepLink integrates heterogeneous datasets that include: i) two genome-wide CRISPR loss-of-function screens: the Broad and Sanger Cancer Dependency Map (DepMap) projects (*n* = 1,054 and 318 cell lines) [1, 2], each covering 17,078 genes; ii) two high-throughput pharmacologic screens: Profiling Relative Inhibition Simultaneously in Mixtures (PRISM primary screen; 4,686 compounds and *n* = 568) [4] and Genomics of Drug Sensitivity in Cancer (GDSC; 287 compounds and *n* = 973) [3], where 205 drugs were shared in common; and iii) the next-generation connectivity map of perturbation signatures called the Library of Integrated Network-based Cellular Signatures (LINCS) [7], which contains changes in expression levels of 12,328 genes perturbed by treatment of 19,811 drugs. Four complementary and interconnected modules in DepLink were designed for different query needs (Fig. 1B). Modules “**1. Query Gene**” and “**3. Query Drug**” take a gene or drug as input and search for perturbations with similar inhibitory effects on cell viability across pan-cancer cell lines or in a specific cancer type. “**2. Query Gene List**” identifies a drug that perturbs the list of target genes by searching among the perturbation signatures of LINCS. Module “**4. Query New Drug**” converts a novel Simplified Molecular Input Line Entry System (SMILES) code to biochemical fingerprints, and then searches for similar drugs. Detailed descriptions of datasets and methods are in the Methods section.

**Fig. 1.**
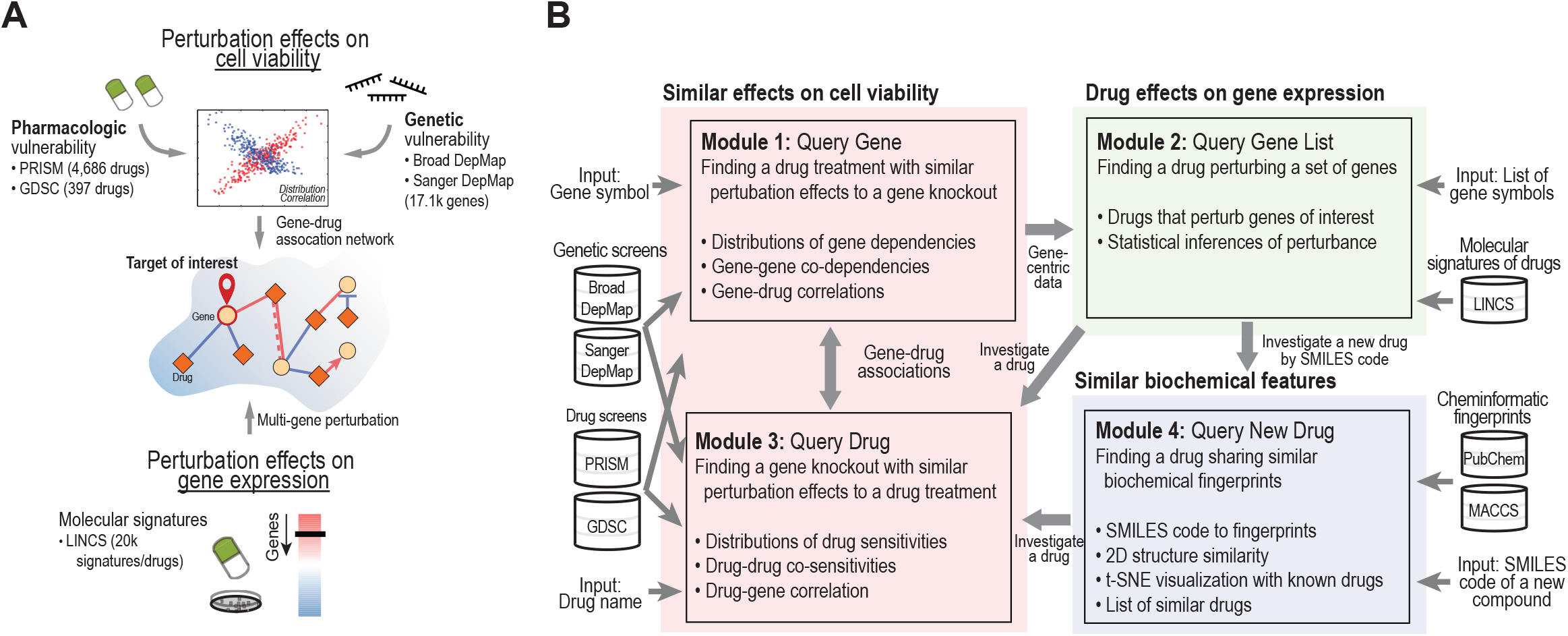
Overview of DepLink. **(A)** Graphic abstract of DepLink. DepLink is an R Shiny app designed to answer two important questions: i) which drugs can be potential surrogates to knockout of a gene in cancer cells, and ii) which genes are potential targets or mechanisms of action of a drug. **(B)** Schematic overview of DepLink. DepLink allows the identification of similar (or opposite) effects induced by genetic and pharmacologic perturbations in cell viability (highlighted in red) and gene expression (green). It systematically incorporates i) two genome-wide CRISPR loss-of-function screens (Broad and Sanger Cancer Dependency Map [DepMap] projects), ii) high-throughput pharmacologic screens (Profiling Relative Inhibition Simultaneously in Mixtures [PRISM] and Genomics of Drug Sensitivity in Cancer [GDSC]), and iii) a next-generation Connectivity Map of perturbation signatures (Library of Integrated Network-based Cellular Signatures [LINCS]). DepLink also allows a user to investigate a new drug of interest based on its biochemical features (highlighted in indigo). Abbreviations: MACCS, Molecular ACCess System; SMILES, Simplified Molecular Input Line Entry System.

We started by building and investigating a global correlation network among all gene knockouts and drugs. Gene-drug, gene-gene, and drug-drug correlation coefficients were calculated based on 456 common cell lines between the Broad DepMap (Chronos gene-effect scores that infer gene fitness effects; see Methods) and PRISM (log-fold changes in cell counts associated with drug treatments) datasets. To generate the network, 2,879 genes with known targets from the PRISM, COSMIC [8], PharmGKB [9, 10], and DrugBank [11] databases that were present in Broad DepMap were selected. A network of 1,019 edges among 449 nodes (291 drugs and 158 genes) was constructed using criteria that yielded manageable numbers of gene-drug (441 pairs), gene-gene (102), and drug-drug pairs (476) (see Methods). In the network, we observed clustering among drugs that were designed to inhibit important oncogenic and cancer progression pathways, such as MEK inhibitors (*e*.*g*., selumetinib and cobimetinib) with genes in the BRAF-MAPK pathway (*e*.*g*., *BRAF, MAPK1*, and *MAP2K1*) and inhibitors of the ErbB family (gefitinib, osimertinib, and dacomitinib). Genes involved in the mitochondrial respiratory chain formed a dense cluster that included cytochrome c oxidase genes (*COX5B, COX6B*, and *COX7C*) and NADH:ubiquinone enzymes (*NDUFA2, NDUFS1*, and *NDUFS2*). *MDM2* inhibitors (nutlin-3 and idasanutlin) formed a cluster with many DNA repair genes that govern cell apoptosis (*TP53, CHEK2, ATM*, and *MDM2*). Three important players in the cell cycle, *CDK6, CCND1*, and *CCNE1*, were connected to palbociclib, which was the first CDK4/6 inhibitor approved for cancer therapy. The results are available under the “**Global Network**” tab of DepLink.

As a use case of DepLink, we further investigated *CDK6*, since it is one of the most promising targets of cell cycle-based cancer therapy [12]. Here we summarize results from multiple modules and focus our analyses on the Broad DepMap and PRISM screens to study similarities in cell viability inhibition induced by *CDK6* knockout, other gene knockouts, CDK4/6 inhibitors, and other drugs; and explore new inhibitors of *CDK6. CDK6* is an essential gene in 109 of 1,054 cell lines assayed by the Broad DepMap. In particular, viability of myeloma, leukemia, and neuroblastoma cells strongly depended on *CDK6* (median scores, -1.02, -0.90 and -0.78). We calculated a Pearson correlation coefficient between the gene-effect scores of *CDK6* and log-fold changes of cell viability induced by each drug assayed by PRISM across 456 common cell lines. Two clusters were formed among the top 10 drugs achieving similar effects with *CDK6* knockout.

One included two CDK4/6 inhibitors, palbociclib and ribociclib (*ρ* = 0.30 and 0.18), as well as ponatinib (which promotes G1 cell cycle arrest and synergizes with palbociclib) [13, 14]. Palbociclib and ribociclib showed highly similar effects in cell viability (*ρ* = 0.58 across 568 cell lines), well beyond many other known *CDK4* and/or *CDK6* inhibitors.

The other cluster of drugs correlated with *CDK6* knockout was composed of seven MEK inhibitors. Cobimetinib is FDA-approved to treat *BRAF*-mutated metastatic melanoma in combination with vemurafenib. Among all types of cancer, skin cancer cells responded best to cobimetinib (median of log fold-changes in viability, -1.93; *n* = 43). The seven MEK inhibitors formed an interconnected network. All were among the top 10 highly correlated drugs of cobimetinib, with correlation coefficients ranging between 0.88 (PD-0325901) and 0.69 (selumetinib).

Both CDK4/6 and MEK inhibitors inhibit cell proliferation. MEK inhibitors target the MAPK pathway, which is responsible for regulating expression levels of D-type cyclins; these in turn regulate cell cycles. Cyclin D regulates *CDK4* and *CDK6* activation, and therefore plays a crucial role in various cancer types [15, 16]; both *CDK4* and *CDK6* are part of the G1 cell cycle checkpoint and allow cells to enter S-phase [17].

The “**Query New Drug**” module of DepLink allows users to investigate a novel drug by a SMILES code and analyze its proximity to known drugs based on molecular fingerprints. To demonstrate the module, we analyzed SHR6390, an investigational CDK4/6 inhibitor now in phase 2 and 3 clinical trials. Preclinical studies of SHR6390 suggested it had antitumor activity in esophageal squamous cell carcinoma [18] and could overcome resistance to endocrine therapy in ER+ and Her2+ breast cancer [19]. DepLink calculated Tanimoto similarities between SHR6390 and all drugs screened by PRISM based on 1,047 biochemical features included in the molecular fingerprint. Palbociclib and ribociclib were the top and third most similar drugs (Tanimoto similarity = 0.96 and 0.82, respectively, with 237 and 213 common biochemical features with SHR6390). TGX-221, a PI3K inhibitor, was the second most similar drug to SHR6390 (Tanimoto similarity, 0.82).

## Conclusions

DepLink is, as far as we know, the first software package that allows users to navigate, visualize, and link rapidly evolving genetic and pharmacologic dependency maps and perturbation signatures of cancer. With a use case of *CDK6*, we demonstrated the capability of DepLink to identify well-characterized drug-gene relationships, and shed light on synergistic effects between CDK and MEK inhibitors and investigational CDK4/6 inhibitors.

## Methods

### Data sources

We incorporated the two largest CRISPR loss-of-function screens from the Broad (21Q4) and Sanger (21Q2) DepMap projects. For both datasets, we used Chronos-normalized gene-effect scores to measure gene fitness effects [2], where a more negative value indicates strong essentiality of the gene in a cell line. For drug screens, we used i) PRISM primary chemical-perturbation viability screening (version, 19Q4), which measures treatment response by log-fold changes in cell counts relative to DMSO; and ii) GDSC (version 2019), based on the half-maximal inhibitory concentration (IC_50_). All screening datasets were downloaded from the Broad DepMap Data Portal (https://depmap.org/portal/download). We also downloaded molecular signatures of drugs from the LINCS database [20] and reanalyzed the data to quantify the effects of a drug on multiple genes for Module 2.

### Analyses of similarities among gene knockouts and drug treatments

The main goal of DepLink is to link CRISPR and drug screens to identify similar inhibitory effects on cell viabilities between a gene knockout and a drug treatment. Pearson correlation coefficients were computed between gene-effect scores (similarities between two gene knockouts), between gene-effect scores and drug responses (between a knockout and a drug treatments), or between drug responses. We constructed a global correlation network using pairwise analysis among all genes from the Broad DepMap data and all drugs in the PRISM screen using 456 common cell lines between the two datasets. Criteria were set at correlation coefficients of 0.65, 0.65, and 0.25 for gene-gene, drug-drug, and gene-drug pairs, respectively. We chose these criteria to achieve manageable numbers of edges among the three groups. In the DepLink tool, users can specify different criteria using slide bars. Pairs meeting the criteria are merged in a network with nodes representing genes and drugs, and edges denoting correlations. For a queried gene or drug, its correlation coefficients with any genes and drugs are tabulated, and the top 10 results are visualized in a network. See the help pages at DepLink for more methods regarding data visualization, identification of drugs perturbing a set of genes, biochemical analyses of new drugs, and tool implementation and usage.

## Acknowledgements

The authors thank Yi-Pei Chen for suggestions and assistance on visual aspects of the server interface.

## Authors’ contributions

TN, LJW, YCC, and YC conceived the study and designed the methods. TN, MN, and YCC collected data. TN, LJW, MN, GR, EJ, YCC, and YC performed data analysis. TP, LJW, and MN implemented the Shiny server. TN, YCC, and YC wrote the manuscript with input from all other authors. All authors interpreted the data. All authors read and approved the final manuscript.

## Ethics declarations

### Ethics approval and consent to participate

Not applicable.

## Consent for publication

Not applicable.

## Availability of data and materials

The DepLink server is freely available at https://chenlabgccri.shinyapps.io/DepLink_dev/.

Demonstrating examples and detailed user manual are provided with the server.

